# Detection and decontamination of chronic wasting disease prions during venison processing

**DOI:** 10.1101/2024.07.23.604851

**Authors:** Marissa Milstein, Sarah C. Gresch, Marc D. Schwabenlander, Manci Li, Jason C. Bartz, Damani N. Bryant, Peter R. Christenson, Laramie L. Lindsey, Nicole Lurndahl, Sang-Hyun Oh, Gage R. Rowden, Rachel L. Shoemaker, Tiffany M. Wolf, Peter A. Larsen, Stuart S. Lichtenberg

## Abstract

Prion diseases, including chronic wasting disease (CWD), are caused by prions, which are misfolded aggregates of normal cellular prion protein. Prions possess many characteristics that distinguish them from conventional pathogens, in particular, an extraordinary recalcitrance to inactivation and a propensity to avidly bind to surfaces. In mid to late stages of CWD, prions begin accumulating in cervid muscle tissues. These features collectively create scenarios where occupational hazards arise for workers processing venison and pose risks to consumers through direct prion exposure via ingestion and cross-contamination of food products. In this work, we show that steel and plastic surfaces used in venison processing can be directly contaminated with CWD prions and that cross-contamination of CWD-negative venison can occur from equipment that had previously been used with CWD-positive venison. We also show that several decontaminant solutions (commercial bleach and potassium peroxymonosulfate) are efficacious for prion inactivation on these same surfaces.

## INTRODUCTION

Chronic wasting disease (CWD) is a fatal prion disease affecting cervids caused by a self-templating, misfolded, and infectious form of the prion protein (PrP^Sc^)(1). Since its discovery in the United States in the 1960s (2,3), CWD has been detected in free-ranging and captive cervid populations in 34 US states, five Canadian provinces, as well as Nordic countries and South Korea (4). CWD continues to spread in white-tailed deer (WTD, *Odocoileus virginanus*), mule deer (*Odocoileus hemionus*), moose (*Alces alces*), and elk (*Cervus canadensis*) populations across the United States and Canada. In the United States alone, over six million white-tailed deer are harvested annually, many of which are consumed and represent an important source of protein for communities across the country (5). A 2017 estimate suggests that as many as 15,000 CWD-positive white-tailed deer are consumed in the USA annually (6). This number, however, is likely underestimated given the limitations of existing CWD surveillance programs and venison food donation efforts within CWD-endemic regions.

CWD prions accumulate in tissues during disease pathogenesis and can be detected in the muscle tissue of white-tailed deer (7), thus, the consumption of meat from CWD-positive cervids may expose humans to CWD prions.. There is concern in the scientific and public health communities about the potential transmission of CWD to humans, particularly via ingestion. The US Centers for Disease Control and Prevention (CDC) acknowledge this risk and recommend reducing the risk by testing cervids before consuming the meat, processing each animal individually to avoid cross-contamination, and not consuming CWD-positive meat (8). There have been no confirmed cases of CWD in humans (9); however, as new CWD strains are identified (10) and more organisms are exposed to PrP^Sc^ (11), there is growing concern that the species barrier may be crossed.

When PrP^Sc^ are introduced into the environment through natural shedding (12)(13) or carcass decomposition (14), they can adsorb to surfaces where they can be detected long after deposition (15)(16)(17). Surface swabbing is an effective CWD detection method in both laboratory settings (18) and natural, environmentally exposed surfaces (19). The study by Yuan et al. (18) highlights the importance of surface structure for prion recovery, noting that porous surfaces, such as wood, were ineffective for swab-based detection, as opposed to non-porous surfaces, such as glass and stainless steel. These attributes (e.g., environmental stability, swab detection, surface adsorption) also factor into surface decontamination, as chemical decontaminants must physically contact PrP^Sc^ aggregates for disintegration or other forms of inactivation (20–22).

Venison processing for human consumption, both in-home and commercial, is an area of potential PrP^Sc^ cross-contamination and direct human exposure. Surfaces, tools, and equipment used for venison processing can be contaminated with PrP^Sc^ from CWD-positive venison (23,24). Cleaning strategies used during venison processing can vary widely and may not be effective in removing or destroying PrP^Sc^, particularly in unregulated facilities (e.g., home butchery, seasonal ‘pop-up’ processors). Thus, there is a clear need to understand the potential of PrP^Sc^ to more widely enter the food supply through surface contamination and the efficacy of chemical decontamination.

In this study, we examined prion contamination of commonly used meat-processing equipment, including knives, cutting boards, and household-style meat grinders, and the efficacy of decontaminants commonly used in home and/or commercial processing. Additionally, we investigated the cross-contamination of CWD-negative meat after contacting CWD-contaminated processing equipment.

## METHODS

Refer to the Supplemental Materials section for a full description of our experimental design. There were five distinct phases of this work (Fig. 1).

**Figure 1.**
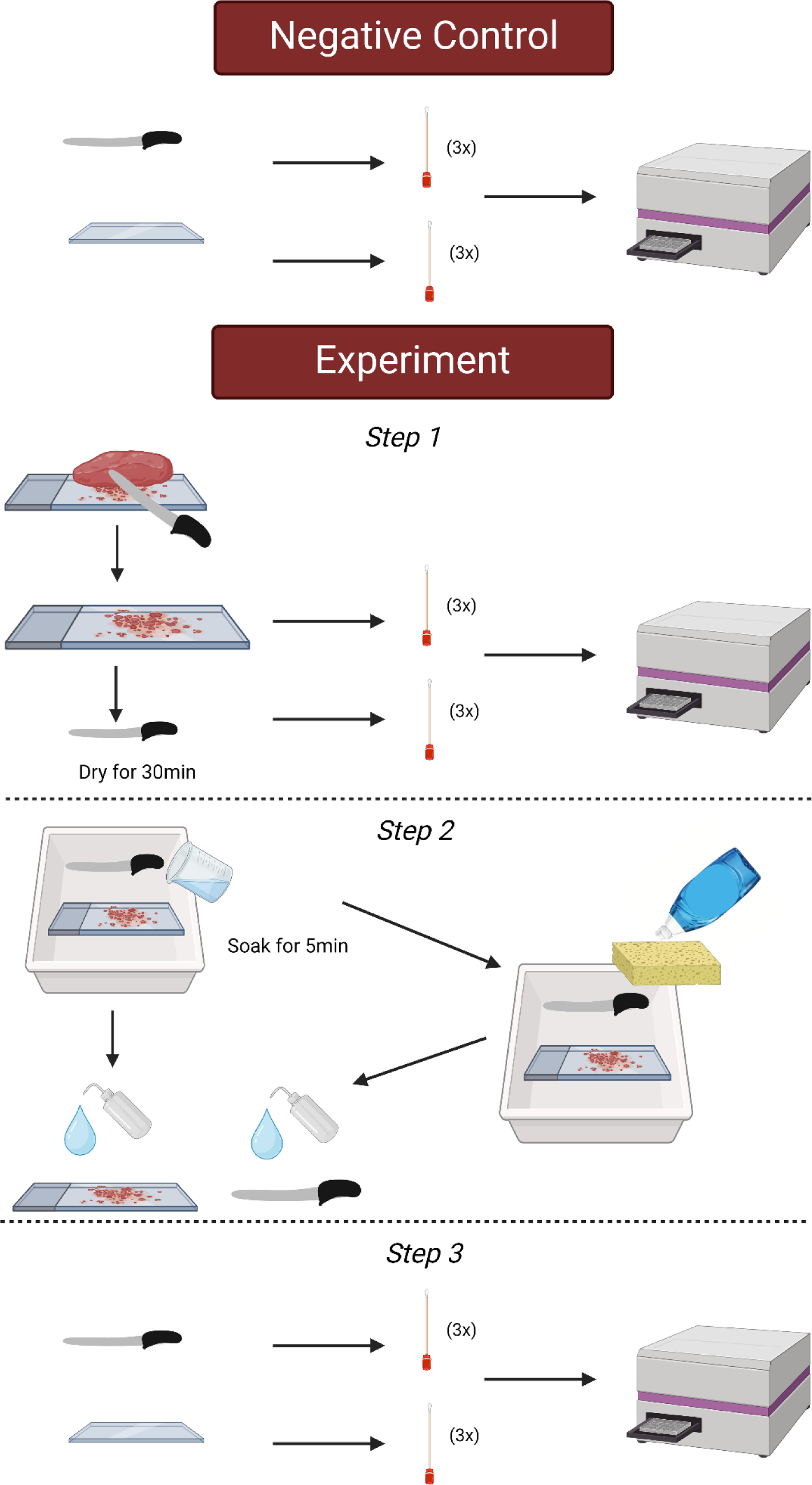
Experimental design of the knife and cutting board study. Negative Control: Surfaces were swabbed before use. Experiment Step 1: CWD-negative or CWD-positive muscle was cut and surfaces were swabbed. Experiment Step 2: surfaces were cleaned. Experiment Step 3: surfaces were swabbed again and swabs were tested by RT-QuIC.

### Pilot Study

We conducted a pilot study to determine whether we could recover, detect, and decontaminate prions on common meat processing surfaces with known CWD-positive samples with different prion loads: 1) WTD cerebellum (high prion load) 2) WTD muscle (low prion load). Control experiments were conducted to determine whether the decontaminants could induce or suppress seeding activity during RT-QuIC. These experiments are detailed in the Supplemental Materials.

### Study Controls

We performed experiments to test if our chosen decontaminants (1) dish soap (Procter and Gamble, https://dawn-dish.com/), 2) Briotech (0.02% hypochlorous acid, Briotech, https://briotechusa.shop), 3) 2% Virkon-S (potassium peroxymonosulfate, Lanxess AG, https://lanxess.com), 4) 10% v/v/ commercial bleach and 5) 40% v/v/ commercial bleach solutions (7.5% sodium hypochlorite; The Clorox Company, https://www.clorox.com)) would interfere with the RT-QuIC assay and if the decontaminants would inhibit prion seeding activity on both stainless steel and cast iron surfaces. In addition, we conducted experiments to assess the recovery and detection of CWD-prions from stainless steel and cast iron surfaces.

### Knife and Cutting Board

We sought to determine whether prion seeding activity could be detected before and after decontamination on stainless steel knives and cutting boards – two standard pieces of equipment used in home and commercial meat processing. We designated a knife and cutting board for each of the five chosen decontaminants. Negative controls were collected by swabbing the knife and cutting board before contact with any muscle samples. To test the efficacy of dish soap for prion decontamination, we made two cuts through CWD-negative muscle on the cutting board. The knife and cutting board were immediately swabbed after the first cut and left to dry at room temperature for 30 minutes after the second cut. A tray was filled with dish soap and water per manufacturer recommendations, where the knife and cutting board were scrubbed with a sponge (3M, https://www.scotch-brite.com) and then rinsed with low-pressure municipal cold water. The surfaces were swabbed again. The experiment was repeated using CWD-positive muscle and new materials. All swabs were stored at −80°C until use.

#### Briotech, 2% Virkon-S, 10% Bleach, 40% Bleach

To test the efficacy of Briotech, 2% Virkon-S, 10% v/v bleach and 40% v/v bleach solutions, we followed the same procedure described above, but no sponge step was performed. Instead, after drying the cutting board and knife for 30 minutes, we soaked the items in the decontaminant solution for five minutes. Then, we rinsed the knife and cutting board with water and swabbed them again. Figure 1 shows a schematic of the knife and cutting board experimental setup.

### Meat cross-contamination

CWD-positive muscle samples were passed through a meat grinder several times to produce a homogenized pool of CWD-positive muscle, which was then sub-sampled. This pool of CWD-positive material was used for the meat grinder experiments. The grinder was disassembled, the gross material was removed, and the parts were swabbed. The grinder was reassembled, and CWD-negative muscle was passed through the grinder and sub-sampled.

### Meat grinder

Based on preliminary results demonstrating the efficacy of the decontaminants used in the knife and cutting board study, we chose the following four decontaminants to be used in experiments using meat grinders: 1) dish soap, 2) Virkon-S, 3) 10% bleach, and 4) 40% bleach solutions. Stainless steel (CHOLISM, amazon.com) and cast iron (CucinaPro, amazon.com) meat grinders were used with each decontaminant. Each grinder was disassembled, and the worm spindle, plate cutter holes, and screw ring threads were swabbed before contact with CWD-positive muscle. These experiments were designed to mimic home or small-scale commercial meat processing.

#### Dish soap - Cleaning of grinder after CWD-positive homogenate

Each grinder was assembled, CWD-positive homogenate was passed through, and allowed to air dry for 30 minutes. The grinders were disassembled, gross material was removed, and the grinder parts were swabbed again. The parts were then placed in a tray with diluted dish soap, and a sponge was used to wash them before we rinsed them with water. We sampled the sponge and swabbed the grinder parts.

#### Virkon-S, 10% Bleach, 40% bleach - Cleaning of grinder after CWD-positive homogenate

For the remaining three decontaminants (Virkon-S, 10% bleach, and 40% bleach), we left the grinder parts to soak for 5 minutes (instead of washing with a sponge), rinsed them with low-pressure cold water, and swabbed in the same locations. Figure 2 shows a schematic of the grinder experimental setup.

**Figure 2.**
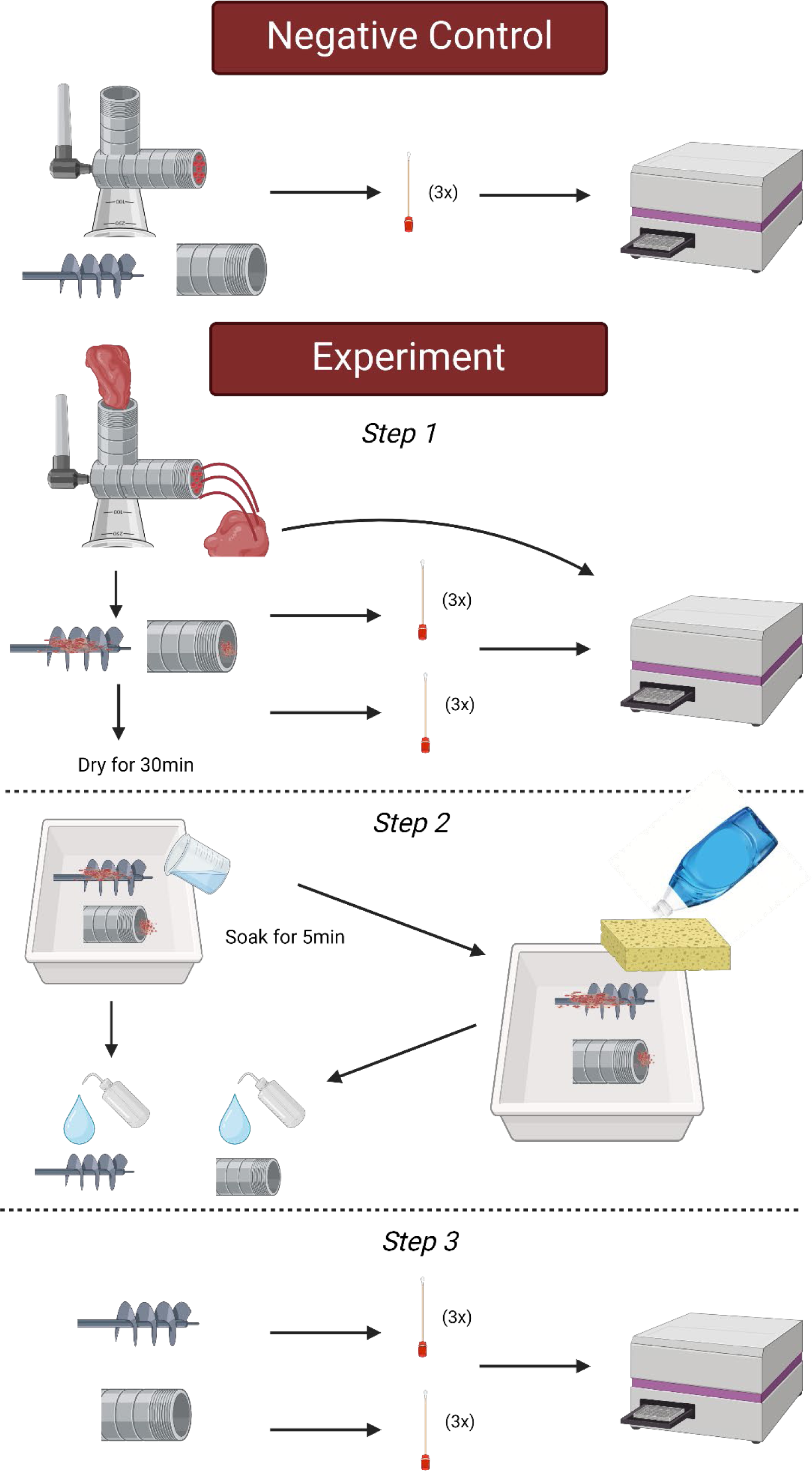
Experimental setup of the meat grinder study. Negative Control: Surfaces were swabbed before use. Experiment Step 1: CWD-negative or CWD-positive muscle was passed through the grinder and surfaces swabbed. Experiment Step 2: grinders were disassembled and surfaces were cleaned/decontaminated. Experiment Step 3: grinder surfaces were swabbed again, and swabs were tested by RT-QuIC.

### Swab Extraction and Processing

Swab extraction and processing followed the procedure of Yuan et al. (19), with some modifications, described in the Supplemental Materials.

### Analysis

Fluorescence readout from the real-time quaking-induced conversion (RT-QuIC) assay captures the kinetics of amyloid formation *in vitro* (25). Three metrics: 1) rate of amyloid formation (RAF), 2) maximum slope (MS) (26,27), and 3) maxpoint ratio (MPR) (28), can be used to describe the three stages of amyloid formation in RT-QuIC – nucleation, elongation, and equilibrium, respectively. RAF was calculated per well as the reciprocal of the time required for fluorescence to reach a threshold of twice the background fluorescence. If a well did not reach the threshold, it was assigned a value of zero. Background fluorescence was calculated as the average fluorescent reading per well at cycles two and three to control for any RFU variability amongst wells and to compensate for the greater ThT fluorescence typically found in the first cycle due to viscosity effects. The slope was calculated as the difference between the relative fluorescence units (RFU) at the current time position plus 6 cycles (3.75 hours) divided by the time. MPR was calculated as the maximum RFU divided by the background RFU. At all three stages, genuine amyloid formation of rPrP induced by CWD prions in RT-QuIC reactions should generate fluorescent curves that are significantly different from reactions with no seeding activities.

A member of the research team (ML), blinded to sample identity and treatment, analyzed the data generated from the pilot study. Using a one-tailed Wilcoxon Rank Sum test, with an ɑ-level of 0.05, RAF, MPR, and MS were compared against the negative plate controls. ML and unblinded team members (MM, SG) reviewed the results from the pilot data and set the following swab inclusion criteria for the remainder of the study: a) all three metrics (RAF, MS, MPR) must be statistically significantly higher than the negative plate controls to be considered positive, and b) regardless of dilution level, if a sample meets criteria (a), the sample will be considered positive. All statistical analyses for swab samples were performed in R (www.R-project.org).

For all muscle sample analyses, we used an uncorrected Fisher’s LSD test with a single pooled variance (ɑ-level of 0.05). We compared RAF, MPR, and MS against the negative plate controls. All three metrics (RAF, MS, MPR) must be statistically significantly higher than the negative plate controls to be considered positive. If a sample meets these criteria, it is considered positive regardless of the dilution level. All statistical analyses of muscle samples were performed using GraphPad Prism (version 10.0.2; graphpad.com).

## RESULTS

### Pilot Study

Multiple swabs demonstrated significant seeding activity in the RT-QuIC assay: 1) after cutting CWD-positive muscle, knife blade swabs and cutting board swabs; and 2) after cutting CWD-positive brain, knife blade swabs and cutting board swabs. After soaking the knife and cutting board with three dilutions of household bleach (i.e., 10%, 40%, 100%), all post-treatment swabs (n=30; two or three knife swabs and three cutting board swabs per dilution, per sample type) were negative by RT-QuIC.

### Study Controls

All negative controls (both knife and cutting board) were negative by RT-QuIC for each of the five decontaminants following treatment. Table 2. shows the results of decontaminant testing and spiking experiments. For each of the five decontaminants (dish soap, Briotech, Virkon-S, 10% bleach, 40% bleach), all samples had no significant seeding activity when CWD-negative muscle tissue was used during knife and cutting board experiments.

**Table 1.** Overview of the study phases and their goals.

| Study Phase | Goal |
| --- | --- |
| Pilot study | Determine whether we could recover, detect, and decontaminate prions on common meat processing surfaces. |
| Study controls | Confirm that the selected decontaminants did not interfere with the RT-QuIC assay, and that CWD-positive material could be detected on both stainless steel and cast iron surfaces. Determine if the decontaminants inhibit seeding activity. |
| Knife and cutting board | Determine whether prion seeding activity could be detected before and after decontaminating stainless steel knives and cutting boards. Test the efficacy of 5 decontaminants on these surfaces. |
| Meat cross-contamination | Determine if CWD-negative meat becomes CWD-positive after passing through a meat grinder that just processed CWD-positive meat. |
| Meat grinder | Determine whether prion seeding activity could be detected before and after decontaminating stainless steel and cast iron meat grinders. Test the efficacy of 4 decontaminants on the grinders. |

**Table 2.** The selected decontaminants were tested to determine if they directly interfered with the RT-QuIC assay. CWD-positive homogenate was added to surfaces actively being treated by, and after rinsing away the decontaminants to assess if they inhibited prion seeding activity. P-values (alpha = 0.05) are shown for RAF (rate of amyloid formation), MPR (maxpoint ratio), and MS (maximum slope). Sample replicates found to have statistically significant seeding activity are shaded in red.

|  | Unspiked decontaminant control samples |  |  |  | Decontaminant control samples spiked with CWD-positive homogenate |  |  |  | Decontaminants rinsed then spiked with CWD-positive homogenate |  |  |  |
| --- | --- | --- | --- | --- | --- | --- | --- | --- | --- | --- | --- | --- |
|  | # Positive replicates | RAF | MPR | MS | # Positive replicates | RAF | MPR | MS | # Positive replicates | RAF | MPR | MS |
|  |  | P-value |  |  |  | P-value |  |  |  | P-value |  |  |
| dish soap | 0/3 | 1.0000 | 0.0048 | 0.5000 | 3/3 | 0.0029 | 0.0048 | 0.0048 | 3/3 | 0.0029 | 0.0048 | 0.0048 |
|  |  | 0.0047 | 0.0048 | 0.0857 |  | 0.0029 | 0.0048 | 0.0048 |  | 0.0029 | 0.0048 | 0.0070 |
|  |  | 1.0000 | 0.6191 | 0.7619 |  | 0.0029 | 0.0048 | 0.0048 |  | 0.0029 | 0.0048 | 0.0048 |
| Briotech | 0/3 | 0.5000 | 0.0857 | 0.0857 | 3/3 | 0.0029 | 0.0070 | 0.0070 | 3/3 | 0.0029 | 0.0048 | 0.0048 |
|  |  | 0.3593 | 0.1762 | 0.1762 |  | 0.0029 | 0.0070 | 0.0070 |  | 0.0029 | 0.0048 | 0.0048 |
|  |  | 0.9146 | 0.5429 | 0.5426 |  | 0.0025 | 0.0048 | 0.0048 |  | 0.0029 | 0.0048 | 0.0048 |
| Virkon-S | 0/3 | 0.5000 | 0.4571 | 0.3048 | 1/3 | 0.0126 | 0.0048 | 0.0048 | 3/3 | 0.0029 | 0.0048 | 0.0048 |
|  |  | 0.5000 | 0.0333 | 0.0571 |  | 0.0471 | 0.0571 | 0.1962 |  | 0.0029 | 0.0048 | 0.0048 |
|  |  | 0.8463 | 0.1286 | 0.1286 |  | 0.1537 | 0.1286 | 0.7619 |  | 0.0029 | 0.0048 | 0.0048 |
| 10% bleach | 0/3 | 1.0000 | 0.8714 | 0.9429 | 0/3 | 0.1537 | 0.3810 | 0.5847 | 3/3 | 0.0029 | 0.0048 | 0.0048 |
|  |  | 1.0000 | 0.5429 | 0.9655 |  | 1.0000 | 0.3810 | 0.9879 |  | 0.0028 | 0.0070 | 0.0070 |
|  |  | 1.0000 | 0.0333 | 0.6191 |  | 1.0000 | 0.6191 | 0.9328 |  | 0.0029 | 0.0070 | 0.0070 |
| 40% bleach | 0/3 | 0.5522 | 0.5429 | 0.8038 | 0/3 | 1.0000 | 0.0070 | 0.9963 | 3/3 | 0.0029 | 0.0070 | 0.0070 |
|  |  | 0.5522 | 0.6952 | 0.8714 |  | 1.0000 | 0.0070 | 0.9963 |  | 0.0029 | 0.0070 | 0.0070 |
|  |  | 0.5522 | 0.3048 | 0.7619 |  | 1.0000 | 0.0346 | 0.97890 |  | 0.0029 | 0.0070 | 0.0070 |

**Table 3.** A high-level overview of the study findings.

| Study phase | Result |
| --- | --- |
| Pilot study | Prions can be recovered and detected on common meat processing items. 10%, 40%, and 100% bleach were effective at decontaminating meat processing surfaces. |
| Study controls | The selected decontaminants do not interfere with the assay. Prions could be recovered and detected on surfaces that had processed CWD-positive meat. Surfaces swabbed after processing CWD-negative meat tested negative for seeding activity. |
| Knife and cutting board | Dish soap removed seeding activity from knife blades but not cutting boards. Briotech removed seeding activity from knives but only partially from cutting boards. Virkon-S, 10% bleach, and 40% bleach removed seeding activity from knives and cutting boards. |
| Meat cross-contamination | CWD-negative meat was contaminated (5 of 8 samples tested positive) after passing through a grinder that had processed CWD-positive meat ~5 minutes earlier. |
| Meat grinder | Dish soap removed seeding activity from cast iron grinders but not stainless steel. Virkon-S reduced but did not eliminate seeding activity from cast iron and stainless steel grinders. 10% and 40% bleach removed seeding activity from cast iron and stainless steel grinders. |

**Table 2**. The selected decontaminants were tested to determine if they directly interfered with the RT-QuIC assay. CWD-positive homogenate was added to surfaces actively being treated by, and after rinsing away the decontaminants to assess if they inhibited prion seeding activity. P-values (alpha = 0.05) are shown for RAF (rate of amyloid formation), MPR (maxpoint ratio), and MS (maximum slope). Sample replicates found to have statistically significant seeding activity are shaded in red.

### Knife and Cutting Board

#### Dish Soap

Significant seeding activity was detected on the stainless steel knife and cutting board after cutting CWD-positive muscle (Fig. 3). It was also detected on the cutting board after cleaning with dish soap but not on the knife or sponge after cleaning with dish soap (Fig. 3; sponge data not shown).

**Figure 3.**
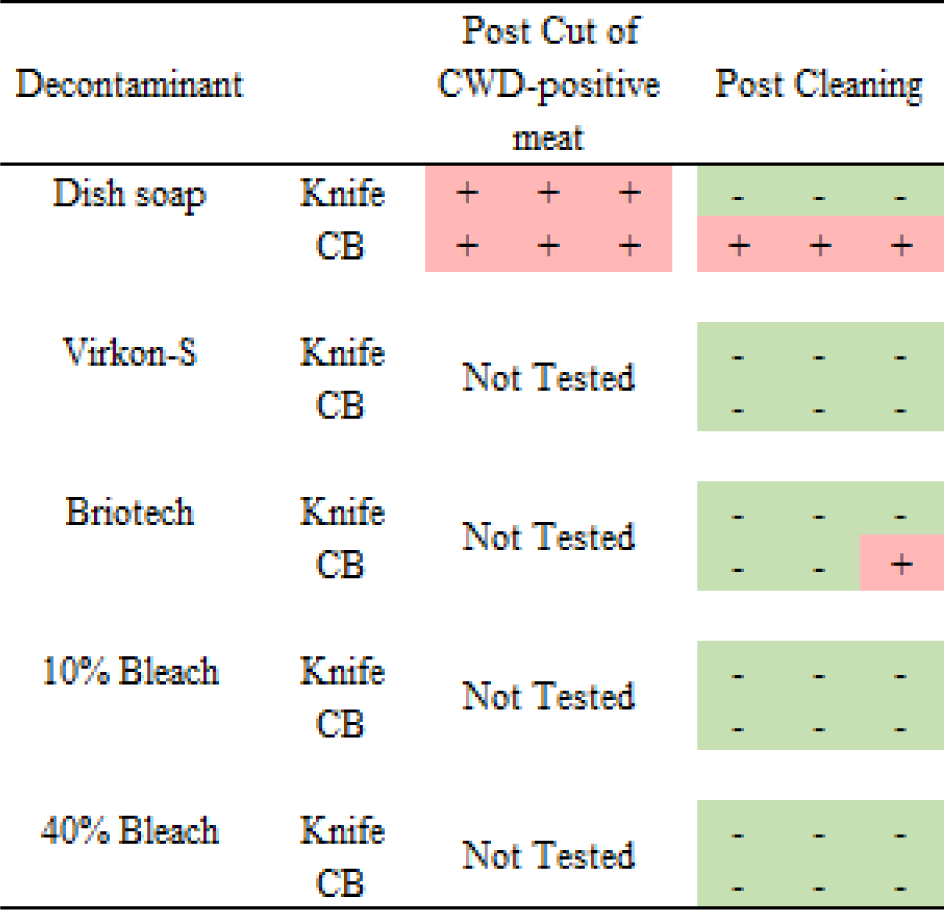
Results from knife and cutting board experiments post CWD-positive muscle cutting and after cleaning with 5 decontaminants (dish soap, Virkon-S, Briotech, 10% bleach, and 40% bleach solutions).

#### Briotech, 2% Virkon-S, 10% Bleach, 40% Bleach

After cleaning the CWD-contaminated cutting board with Briotech, we detected significant seeding activity in one sample (Fig. 3). No significant seeding activity was detected on CWD-contaminated knives after cleaning with any of the decontaminants (Fig. 3). No significant seeding activity was detected on CWD-contaminated cutting boards after decontamination with Virkon-S, 10% Bleach, or 40% Bleach (Figure 3).

### Meat Cross-Contamination

An 88% significant seeding rate was detected in the CWD-positive muscle homogenate pool (7/8 biological replicates; Fig. 4). When CWD-positive muscle homogenate was passed through a grinder, we detected significant seeding activity on the plate cutter (2/3, 66.7%) and screw ring thread (1/3, 33.3%). No seeding activity was detected on the worm spindle of the grinder. When CWD-negative muscle (ID=2106) was passed through the grinder after the CWD-positive muscle homogenate, we found a 63% seeding activity rate (5/8; Fig. 4).

**Figure 4.**
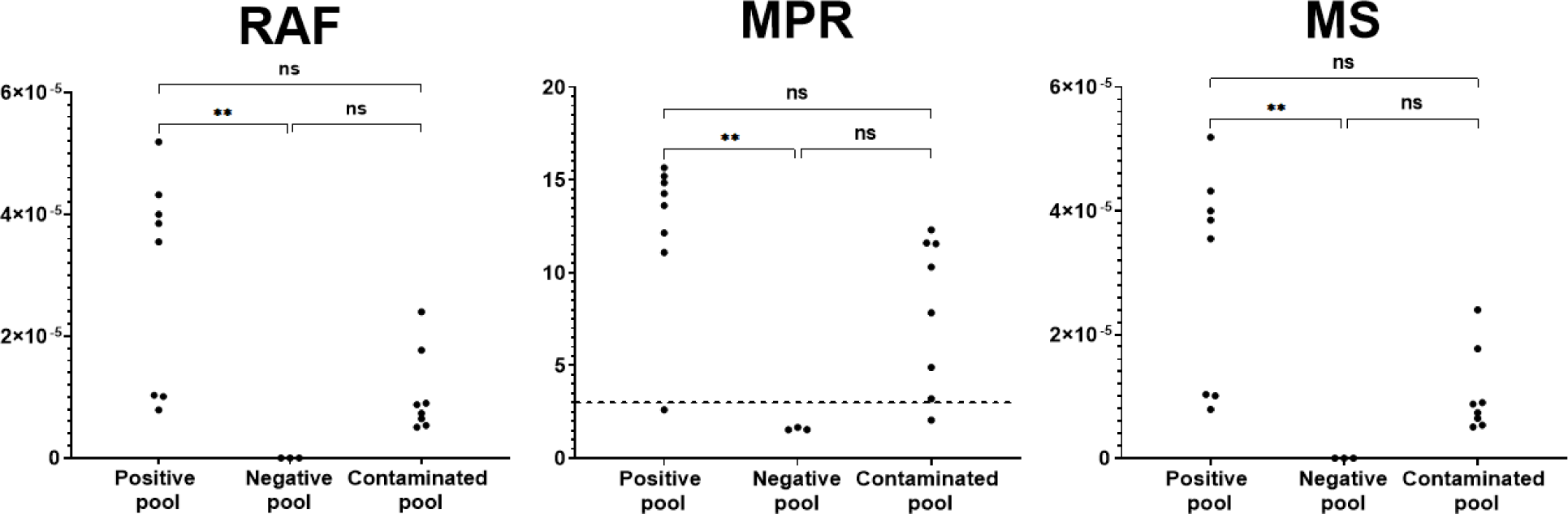
RT-QuIC results from the CWD-positive muscle homogenate (Positive pool), the CWD-negative muscle homogenate before passing through a contaminated grinder (Negative pool), and the CWD-negative muscle homogenate after passing through a contaminated meat grinder (Contaminated pool). RAF=rate of amyloid formation; MPR=maxpoint ratio; MS=maximum slope; ns=not statistically significant (alpha = 0.05); ** indicates a P-value < 0.045.

### Meat Grinder

All negative controls of the stainless steel and cast iron grinder parts were negative by RT-QuIC (Figure 5). Significant seeding activity was demonstrated from swabs from all stainless steel and cast iron grinder parts after passage of CWD-positive muscle homogenate (Figure 5, 7). Significant seeding activity was demonstrated from multiple swabs after dish soap and Virkon-S decontamination of the stainless steel and cast iron grinder parts, while no significant seeding activity was demonstrated from swabs after 10% and 40% bleach decontamination (Figure 6, 7).

**Figure 5.**
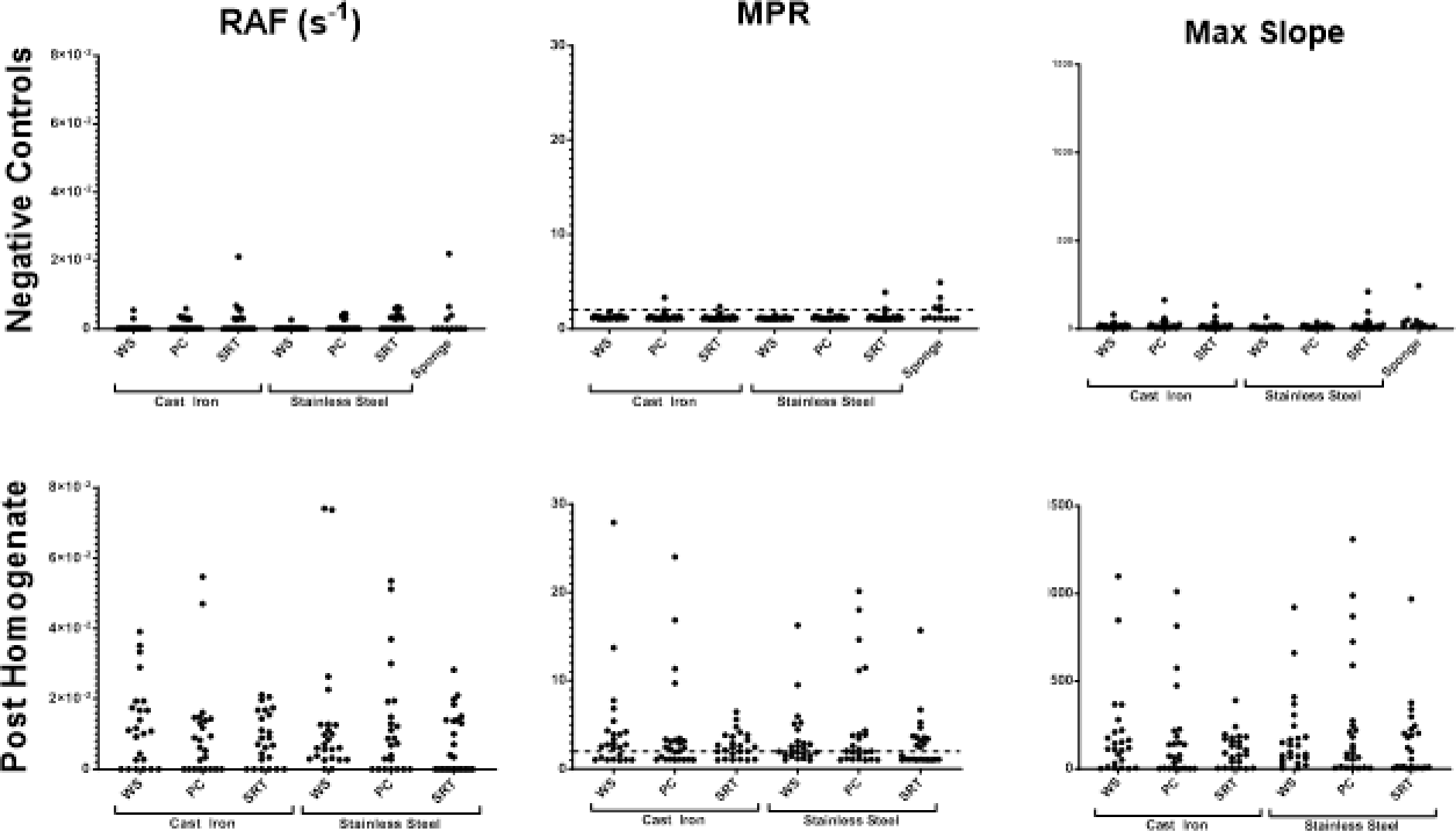
Meat grinder RT-QuIC swab results for negative controls (before use) and after CWD-positive muscle (Post Homogenate) was passed through. Each dot is an average of a single biological replicate (consisting of 8 technical replicates). Each set of paired samples (e.g., cast iron worm spindles, etc.) resulted in a statistically significant difference post homogenate as compared to the negative samples (alpha = 0.05; data not shown). RAF=rate of amyloid formation; MPR=maxpoint ratio; MS=maximum slope; WS=worm spindle, PC=plate cutter, SRT=screw ring threads

**Figure 6.**
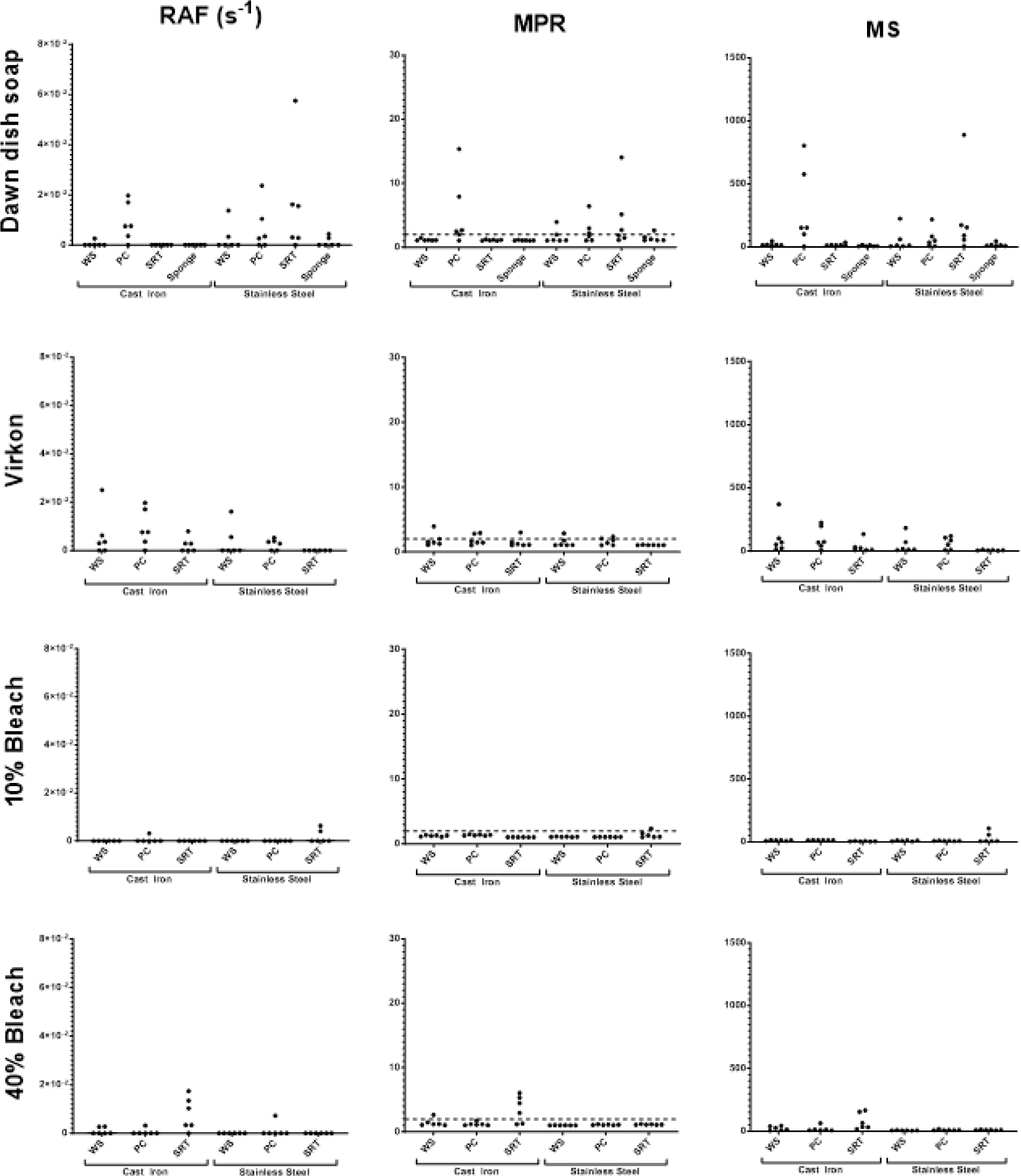
Meat grinder swab results after soaking in one of four decontaminants (after CWD-positive muscle was passed through each). Each dot is an average of a single biological replicate (consisting of 8 technical replicates). RAF=rate of amyloid formation; MPR=maxpoint ratio; MS=maximum slope; WS=worm spindle, PC=plate cutter, SRT=screw ring threads

**Figure 7.**
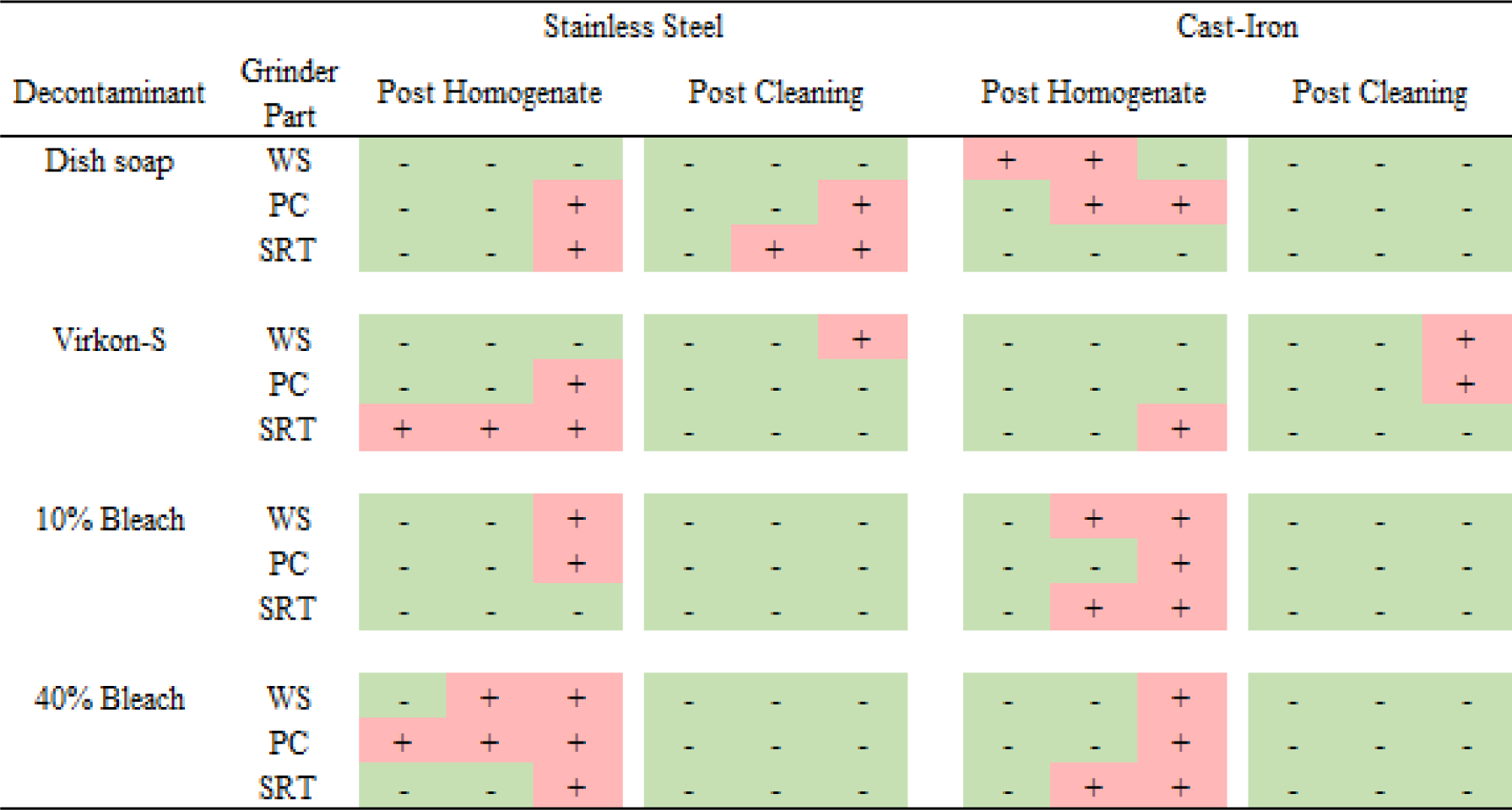
Results from stainless steel and cast-iron grinder experiments post CWD-positive muscle homogenate passing through each grinder and after cleaning with each of the decontaminants (dish soap, Virkon-S, 10% bleach, and 40% bleach solutions). Grinder part abbreviations are as follows: WS= worm spindle, PC= plate cutter, SRT= screw ring threads.

## DISCUSSION

In this study, we examined the contamination of commonly used meat-processing equipment with CWD prions from CWD-positive muscle and the efficacy of decontaminants commonly used in home and/or previously shown to have variable levels of efficacy for prion decontamination (20–22). Additionally, we investigated the cross-contamination of meat in CWD prion-contaminated meat processing equipment. Using a conservative approach in determining whether samples were positive (by requiring positivity using all three RT-QuIC metrics), we found that 1) CWD prions can be detected on common meat processing surfaces after coming into contact with CWD-positive white-tailed deer muscle; 2) CWD prions can be transferred to CWD-negative meat after passing through a contaminated meat grinder; 3) Virkon S, 10% bleach, and 40% bleach were highly effective for CWD prion decontamination on surfaces; and 4) surface composition may play a role in CWD prion detection and decontamination on meat processing equipment.

Based upon our results, it is clear that CWD prion seeding activity is detectable on common meat processing equipment such as knives, cutting boards, and multiple parts of meat grinders. We would note that the presence of seeding activity does not directly imply infectivity, and the relationship between seeding activity and infectivity can be complicated. However, given that seeding activity is correlated to infectivity (29), and we are using materials that have verification of PrP^Sc^ presence by other means, it is reasonable to conclude that the presence or absence of seeding activity in our study can be correlated with the presence or absence of infectivity. Further investigation is warranted into the contamination regimes and decontaminants described here using animal bioassays.

Importantly, this was demonstrated not only with tissues containing high levels of prions (i.e. brain from a CWD-positive animal) but also in muscles with progressively lower levels of prions from a clinical deer and two hunter-harvested deer. We also detected seeding activity in the CWD-negative muscle homogenate after it passed through the contaminated meat grinder. These exemplify “real life” scenarios with implications for the food supply where surfaces and tools are re-used between deer, and meat is mixed from multiple deer, potentially with non-clinical, un-tested deer.

We found that 10% and 40% bleach were highly effective, and Virkon-S to a lesser extent, at decontaminating the meat processing surfaces tested in this study. Briotech was less effective and inconsistent in decontamination. Although the overall findings are promising, further investigation is warranted. It is possible that reduced concentrations of bleach and contact times could still result in effective decontamination, as we did not test the lowest effective duration or concentration of decontaminants, nor did we test how repeated disinfection with these solutions impacts surface integrity and ongoing disinfection efficacy. This is important as these chemicals can compromise the integrity of surfaces such as stainless steel, leading to unknown consequences of prion adsorption and decontamination. Additionally, though we observed a few negative control swabs of grinder parts that were positive on initial test but negative upon retest, we hypothesize that these interferences arose from grease or metallic debris from the machining of parts. From a cleaning protocol perspective, the cleaning/scrubbing of tools, parts, and surfaces is an important step, as tissue debris tends to remain on some surfaces after a decontamination soak. As demonstrated by others (21), decontaminants are ineffective at penetrating tissue, so removal of the tissue remnants will lead to more effective decontamination and reduce cross-contamination. This is one plausible explanation for the reduced efficacy of Virkon-S in one of the aforementioned treatments. The containment and disposal of that pre-decontamination “cleaning wash” is also an important consideration for environmental prion contamination.

Interestingly, we saw differences in the detection of prion seeding activity between the knife and cutting board, where after decontamination with dish soap, we were able to detect prion seeding on the surface of the cutting board but not on the surface of the knife. Curiously, we did not detect prion seeding activity in any of the sponge samples. We cannot discount the possibility that prions were removed from the knife surface by dish soap and simply remained in the decontaminant solution, speaking back to our point regarding containment and disposal of the cleaning solution itself. The principal components of dish soap are surfactants (30), which can remove adsorbed prions from surfaces, resulting in a lack of detection on the steel surface, while also diluting the prions to the point that they are undetectable by RT-QuIC. These factors must all be considered when planning and implementing cleaning and decontamination of environmental prions.

In conclusion, our results show that the processing of CWD-positive cervids has the potential to contaminate meat-processing equipment and cross-contaminate downstream meat products. Further, our data and those previously published (20,21) indicate that Virkon-S and bleach solutions with appropriate contact times (as little as five minutes) can effectively decontaminate non-porous surfaces of CWD prions. Following the BSE outbreak in Europe in the early 1990s, processing practices became mandatory to remove and avoid specified risk materials (SRM, e.g., spinal cord, brain) and incorporate downstream molecular testing procedures to identify contamination in meat products (31). Our findings indicate similar practices may be necessary to reduce CWD exposure in humans through meat processing. However, given the unique features of CWD prions, contamination of equipment and surfaces is more challenging to control. This further highlights the importance of testing cervids prior to processing, as well as compliance with effective meat processing, environmental screening, waste management, and cleaning/decontamination protocols.

## BIOGRAPHICAL SKETCH

Dr. Marissa Milstein is a wildlife veterinary researcher at the University of Minnesota in St. Paul. Her primary research interests involve studying zoonotic disease transmission from the hunting, butchery, and consumption of wildlife.

## ACKNOWLEDGEMENTS

We thank Davis Seelig and Russ Mason for their helpful discussions in planning this project. Sam Thomas kindly assisted with preliminary RT-QuIC experiments. We also thank Nicole Neeser from the Minnesota Department of Agriculture for guidance regarding meat processing and regulations. We are grateful to the Minnesota Department of Natural Resources, the University of Minnesota Veterinary Diagnostic Lab, and deer hunters for providing samples. We thank Suzanne Stone for lab management, Adam Reinschmidt for assisting with the pilot study, and Corina Valencia for sample management. Professor Joel Pedersen contributed substantially to the acquisition of funding from the Michigan Department of Natural Resources. This project was also funded by the Minnesota legislature through the Rapid Agricultural Response Fund, the Michigan Department of Natural Resources, the Minnesota Agricultural Research, Education, Extension, and Technology Transfer (AGREETT) program, and the Minnesota Environment and Natural Resources Trust Fund, as recommended by the Legislative-Citizen Commission on Minnesota Resources (LCCMR).

