## Supplemental Materials for "Detection and decontamination of chronic wasting disease prions during venison processing"

Methods

*Positive and Negative Control Samples*

CWD-positive and CWD-negative control tissues were opportunistically collected for disease testing purposes from the Minnesota Department of Natural Resources (MN DNR), Minnesota Board of Animal Health (MN BAH), or by licensed hunters. CWD-positive or negative status was determined by ELISA and/or IHC (Table S1). No animals were specifically sacrificed for this study, and the study was determined to be exempt from Institutional Animal Care and Use Committee (IACUC) approval.

ELISA characterized CWD-positive and CWD-negative brain, lymph node, and muscle samples were tested by RT-QuIC, as described below, to be used as control samples in this study. Our main goals were to determine whether the samples were viable as negative or positive control samples and establish a semi-quantitative baseline of seeding activity. Cerebellum and lymph nodes were tested after diluting to 10^-3^, muscles were tested undiluted, and at 10^-1^.

| **ID** | **AnimalCWD status** | **Age/sex** | **Scientific name** | **Cause of death** | **Muscle collected** | **Test used to determine CWD status** | **Tissues used for CWD testing** |
| --- | --- | --- | --- | --- | --- | --- | --- |
| 297 | Positive | Yearling, male | *Odocoileus virginianus* | Hunter harvested | Shoulder | IHC, ELISA | medial retropharyngeal lymph nodes |
| 307 | Positive | Adult, female | *Odocoileus virginianus* | Farm raised, found deceased | Front leg, rear leg, backstrap, cerebellum | IHC | cerebellum, medial retropharyngeal lymph nodes, prescapular lymph node |
| 31 | Negative | Adult, female | *Odocoileus virginianus* | Farmed raised, euthanized | NA | IHC, ELISA | Parotid lymph node |
| 2106 | Negative | Adult, male | *Odocoileus virginianus* | Hunter harvested | Neck | RT-QuIC | Obex, medial retropharyngeal lymph nodes |
| 2324 | Positive | Adult, female | *Odocoileus hemionus* | Hunter harvested | Tenderloin | IHC, ELISA | medial retropharyngeal lymph nodes |

**Supplemental Table 1.** Control samples used in the study and the test(s) used to determine CWD status. State regulatory agencies performed and reported immunohistochemistry (IHC) and enzyme-linked immunosorbent assay (ELISA) tests. Real-time quaking-induced conversion (RT-QuIC) tests performed in-house.

*Pilot Study*

Decontamination procedures followed the general outline of Sanitation Standard Operating Procedures of the National Science Foundation (32) and the Minnesota Department of Agriculture (33). We selected cleaning products based upon consumer availability and previously demonstrated prion inactivation efficacy (20–22).

We designated one knife and one cutting board for each bleach (6% sodium hypochlorite; The Clorox Company, www.clorox.com) dilution: 1) 10% bleach, 2) 40% bleach, and 3) 100% bleach. Each knife blade (Dexter-Russel, www.dexter1818.com) and cutting board (Crestware, www.crestware.com) was swabbed prior to use to ensure the items were not contaminated or that the materials (e.g., stainless steel knife blade, polyethylene cutting board) would not induce false positive seeding activity in RT-QuIC (negative control). We then placed a positive control tissue (cerebellum, ID =307 and front leg muscle, ID =307 front leg) on the cutting board and cut the tissue, ensuring the entire length of the stainless steel knife blade came into contact with the tissue sample. We swabbed the knife blade and cutting board surface using Fisherbrand^TM^ PurSwab Foam Swabs (Thermo Fisher Scientific, www.fishersci.com). Each swab was taken from an area in contact with tissue, and we did not swab the same area twice. Each surface (i.e., knife blade and cutting board) was set to dry at room temperature for five minutes, rinsed with 10mL of PBS to remove gross debris, and swabbed again with a single swab. The knife and cutting board were then soaked in the designated bleach solution (10%, 40%, 100%) for five minutes, removed, rinsed with tap water, and allowed to drip dry. The knife blades and cutting boards were swabbed again. All swabs were placed in 15mL conical tubes with 500 μL of PBS and frozen at -20°C until processing.

*Pilot Study Swab and Tissue Processing*

Swab processing was performed following Yuan et al. (18), with the following modifications: 1 X PBS was used, sonication was performed in 3 treatment cycles at room temperature, and centrifugation (21,100 x g at 4C for 30 minutes) was used to concentrate the extracts. Swabs were tested at 4 dilutions (undiluted, 10^-1^, 10^-2^, 10^-3^) in RT-QuIC sample buffer. Muscle processing was performed as described in Li et al. (7) using the freeze-thaw method and muscle samples were tested undiluted and at 10^-1^. We followed the procedures outlined in Schwabenlander et al. (34) to test brain and lymph node samples at 10^-3^.

**Study Controls**

Negative and Positive Control Experiments on Stainless Steel

For the negative control experiment, 50ul of decontaminant was pipetted onto a stainless steel plate, allowed to dry for five minutes, and the location of where the decontaminant was placed on the stainless steel plate was swabbed. The plate and decontaminant were rinsed with low-pressure cold water, dried with Kim wipes (Kimberly-Clark Professional, www.kcprofessional.com), and swabbed again. This process was repeated for each decontaminant in replicates of six swabs per condition. To create the negative control aliquots, 100uL of the swab-processing extract was aliquoted into a new tube and served as the negative control. A 90ul aliquot of the same swab-processing extract was pipetted into a new tube, spiked with 10ul of a 10^-3^ dilution of a CWD-positive prescapular lymph node control (ID= 307), and served as the positive control. For both the negative and positive controls, the extracts were centrifuged at 21,100 x g for 30 minutes at 4°C, the supernatant discarded, and the pellets stored at -80°C until RT-QuIC testing.

Negative and Positive Control Experiments os Cast Iron

To determine whether prions could be successfully recovered from a cast iron surface via swabbing methods and whether decontaminants could induce or suppress seeding activity during RT-QuIC, a negative and positive control experiment was performed for each of four decontaminants (dish soap, Virkon-S, 10% bleach, and 40% bleach). Control experiments followed the methods described in the knife and cutting board study, swabbing only the outside of the cast iron grinder surface. An additional positive control was performed by pipetting positive control (ID= 307; WTD prescapular lymph node) directly onto the surface of the cast iron grinder, and six swabs were collected.

To process the negative control swabs, 100uL of the initial 200uL of swab-processing liquid (1 x PBS) was aliquoted into a new tube. A 90ul aliquot of the remaining liquid was pipetted into a new tube, spiked with 10uL of a 10^-3^ dilution of a CWD-positive prescapular lymph node homogenate (ID= 307), and served as the positive control. For both the negative and positive controls, the tubes were centrifuged at 21,100 x g for 30 minutes at 4℃, the supernatant discarded, and the pellets stored at -80°C until RT-QuIC testing. All of the control swabs were tested undiluted and at 10^-1^.

**Knife and Cutting Board Experiments**

Experimental Design

A single knife and cutting board were designated for each of the following decontaminants: 1) dish soap (Procter & Gamble, www.dawn-dish.com), 2) Briotech (Briotech, www.briotechusa.shop), 3) 2% Virkon-S (Lanxess AG, www.lanxess.com), 4) 10% bleach, and 5) 40% bleach solutions (7.5% sodium hypochlorite; The Clorox Company, www.clorox.com). 7.5% sodium hypochlorite was used in this study (compared to 6.0% sodium hypochlorite used in the pilot study) as the concentration of commonly available bleach changed during the study.

On each cutting board, a square grid (consisting of two columns and three rows for a total of six cells) was drawn with a permanent marker to delineate the placement and swabbing locations of muscle tissue. Similarly, on each side of a knife blade, two lines were drawn with a permanent marker to delineate three swabbing locations (a total of six locations; see Supplemental Figure 1). The cutting board and knife were swabbed for negative control sampling on each of the six locations. Swabs were placed in fresh 1.5mL tubes and stored at -80°C until processing. Six swabs were collected per condition, with three used for testing and the remaining three retained and stored at -80°C should the need arise for additional testing.

*Dish Soap*

To test the efficacy of dish soap for prion decontamination, CWD-negative muscle (ID= 2106, neck muscle) was placed on the cutting board grid and cut with a stainless steel knife, ensuring the tissue came in contact with the entire length of the blade. The muscle was removed from the board, and six swabs (one from each grid cell) were collected. Swabs were then collected from each of the six sections along the knife blade. The muscle tissue was placed back on the cutting board grid, and a second cut was made. The muscle was removed, and the knife and cutting board were set aside to dry for 30 minutes. Using new, unused scissors, six pieces were cut from a new, unused Scotch Brite™ sponge (3M, www.scotch-brite.com) and placed individually in 1.5ml tubes (negative controls). A tray was filled with a 5mL of dish soap to 3.8L of water solution, and the knife and cutting board were scrubbed with the sponge and rinsed with low-pressure, cold water. Six pieces were cut from the used sponge and placed individually in 1.5ml tubes. The knife and cutting board were swabbed again. The experiment was repeated using CWD-positive muscle (ID= 307, front leg) and new materials (e.g., cutting board, knife, tray, etc.). All samples were stored at -80°C.

*Briotech, 2% Virkon-S, 10% Bleach, 40% Bleach*

To test the efficacy of Virkon-S, Briotech, and bleach solutions, we followed the same procedure as described for the dish soap, but no sponge step was performed. Rather, after allowing the cutting board and knife to dry for 30 minutes, the items were soaked in the decontaminant solution for five minutes. After soaking, the knife and cutting board were rinsed with low-pressure cold water and swabbed.

Knife and Cutting Board Experiments - Swab and Muscle Processing

All experimental swab and sponge samples were processed via the modified Yuan et al. methods described above in the pilot study section. All sponge and swab samples used in the knife and cutting board study were tested undiluted and at 10^-1^.

Muscle samples were processed by adding 100 mg (+/- 5 mg) of tissue to 900 ul 1X PBS in a pre-filled 2 ml tube with 1.5 mm zirconium beads (BenchMark Scientific, www.benchmarkscientific.com) and homogenized at max speed in a BeadBlaster™ 24-tube homogenizer (BenchMark Scientific, www.benchmarkscientific.com) for a total of 180 seconds, with a 30 second rest period between homogenizing. The bead tubes were then centrifuged at 2.4 x g for three minutes to separate the solid and liquid phases. 500 ul of the liquid phase was removed, placed in a new 1.5 ml centrifuge tube, and then centrifuged at 21,100 x g for 40 minutes at 4°C. The supernatant was removed, and the pellet was stored at -80°C until testing. Muscle samples were tested undiluted and at 10^-1^.

**Meat Grinder Experiment**

Based on preliminary results demonstrating the efficacy of the decontaminants used in the knife and cutting board study, we chose the following four decontaminants to be used in the meat grinder experiments: 1) dish soap, 2) Virkon-S, 3) 10% bleach, and 4) 40% bleach solutions (7.5% sodium hypochlorite). Stainless steel and cast iron meat grinders were used with each decontaminant, and the following experiments were designed to mimic at-home or small-scale commercial meat processing.

Experimental design:

*Creation of a homogenate of known CWD-positive muscle*

CWD-positive muscle samples (ID=307, 297, 2324) were passed through a meat grinder multiple times to create a homogenized pool of CWD-positive muscle (CWD-positive homogenate). The muscle was collected in a weigh boat and sampled (n= 8). The grinder was disassembled, gross material removed, and the worm spindle, plate cutter holes, and screw ring threads were swabbed again (n= 6). The grinder was reassembled, then CWD-negative muscle (ID=2106) was passed through the grinder and sampled (n=8). The grinder used to create the homogenate was then discarded. A new grinder (stainless steel and cast iron) was used for each decontaminant.

*Dish soap - Decontamination of grinder after CWD-positive homogenate*

A stainless steel grinder was disassembled, and the worm spindle, plate cutter holes, and screw ring threads were swabbed (six swabs each) prior to contact with CWD-positive homogenate. The grinder was assembled, and the CWD-positive homogenate was passed through the grinder and collected in a weigh boat. The grinder was allowed to air-dry for 30 minutes, disassembled, gross material removed, and the same grinder parts were swabbed again (six swabs per part). The parts were then placed in a tray with diluted dish soap as described above. Using new, unused scissors, six pieces of a new sponge were collected and the remaining portion of the sponge was used to wash the grinder parts. The parts were rinsed with low-pressure, cold water. Sponge and grinder parts were sampled/swabbed again. The experiment was then repeated with a cast iron grinder.

*Virkon-S, 10% Bleach, 40% bleach*

For the remaining three decontaminants (Virkon-S, 10% bleach, and 40% bleach), grinder parts were left to soak for 5 minutes (instead of washing with a sponge), rinsed with low-pressure, cold water and swabbed in the same locations.

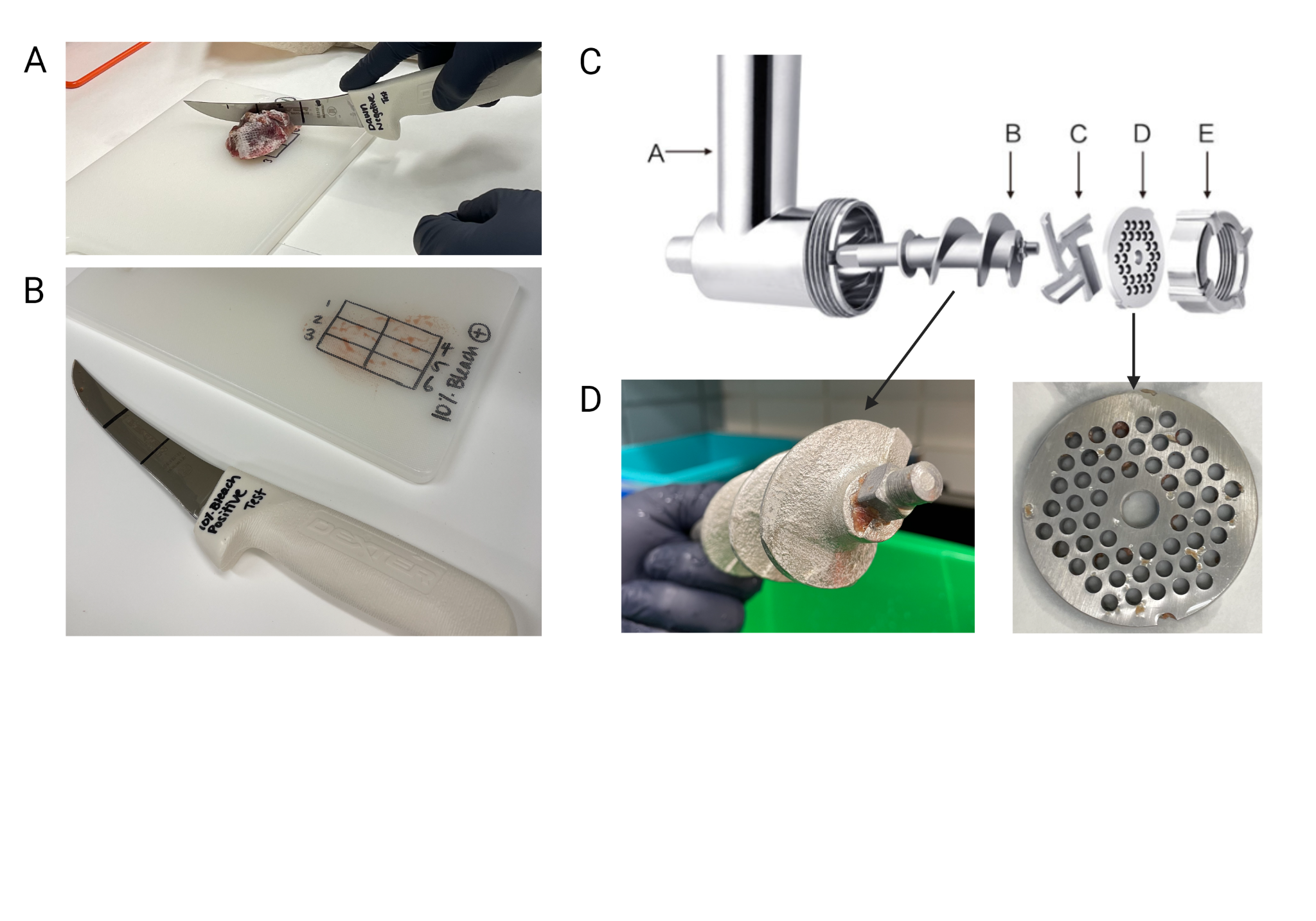

**Supplemental Figure 1.** Parts of the cutting board, knife, stainless steel grinder, and cast iron grinder that were swabbed for prion contamination. Six swabs were taken from each part, ensuring that no area was swabbed more than once. A) Cutting the muscle sample with the knife on the cutting board. B) Cutting board and knife were marked with six locations (three on each side of the knife blade) to ensure areas were not swabbed more than once. C) Schematic of the grinder parts: grinder body (A), worm spindle (B), grinding blade (C), cutting plate (D), screw ring (E). D) Worm spindle (left) and cutting plate post decontaminant soak.

Meat Grinder Experiments - Swab and Muscle Processing:

Swab and muscle samples for the meat grinder study were processed in the same manner as in the knife and cutting board study, as described above.

**RT-QuIC Methods**

RT-QuIC procedures were either performed manually or using an automated liquid handler (Eppendorf epMotion 5070, www.eppendorf.com).

Swab, Brain, Lymph Node, and Pilot Study Muscle

RT-QuIC assay parameters are described by Schwabenlander et al. (34) with the following modifications: intervals of 60 seconds shaking and 60 seconds resting over 48 hours. Each plate contained positive (ID=307, WTD medial retropharyngeal lymph node) and negative (ID=2106, WTD medial retropharyngeal lymph node) assay controls.

Knife/Cutting Board and Meat Grinder Study Muscle

RT-QuIC of the muscle samples used in the knife/cutting board and meat grinder studies were enhanced using silica nanoparticles following the methods of Christenson et al. (2023), hereafter called Nano-QuIC. Nano-QuIC master mixes were made to the following specifications: 1X PBS, 1 mM Ethylenediaminetetraacetic acid (EDTA), 170 mM NaCl, 10 μM thioflavin T (ThT), 0.1 mg/mL rPrP, and 2.5mg/ml 50nm silica nanoparticles (Christenson et al. 2023). The Nano-QuIC instrument parameters were the same as described above.

RT-QuIC for Control and Experimental Swabs:

Each 96-well plate included four technical replicates of a blank (dilution buffer), six replicates of the assay positive control (ID= 307; WTD prescapular lymph node) and six replicates of the assay negative control (ID= 2106; WTD medial retropharyngeal lymph node).

**Supplemental Results**

Positive and Negative Muscle Controls

Muscle samples 307 front leg, rear leg, backstrap; 297 shoulder; and 2324 tenderloin demonstrated significant seeding activity be RT-QuIC, and were from animals with CWD-positive results using retropharyngeal lymph node and/or obex by ELISA and/or IHC (5; Supplemental Table 1; data not shown). Muscle sample 2106 neck and the retropharyngeal lymph node and obex from 2106 demonstrated no seeding activity by RT-QuIC (Supplemental Table 1; data not shown).
